## Supplementary Materials for "Well-Differentiated Papillary Mesothelioma of the Peritoneum is Genetically Distinct from Malignant Mesothelioma"

### Supplementary Text

#### Comparison between Stevers *et al.* 2018 and Shrestha *et al.* 2019 (present study)

Here we provide a detailed comparison between Stevers *et al.* 2018 and the present study which we label Shrestha *et al.* 2019 for convenience.

Stevers *et al.* used a targeted panel of 479 cancer-related genes (UCSF500 Cancer Panel) for sequencing (Illumina HiSeq 2500 machine), whereas we used Ion AmpliSeq™ (Thermo Fisher Scientific) Exome Sequencing which covers 18,961 genes (Ion Proton™). The overlap in the genes examined between these two studies is given in **Supplementary Figure 5A**.

Using a targeted panel provided Stevers *et al.* an advantage to sequence a small number of genes at a high depth (average depth = 320x, range = 33x - 722x), whereas we sequenced a large number of genes at a cost of sequencing depth (average depth = 102x).

Stevers *et al.* reported 21 mutations covering 10 genes in 10 WDPM cases, where as we have identified 461 mutations covering 297 genes in 5 WDPM cases. There is no overlap of the mutated genes reported in Stevers *et al.* and our study (**Supplementary Figure 5B**). In fact, UCSF500 gene panel used by Stevers *et al.* covered only 10 mutated genes reported by our study (**Supplementary Figure 5C**).

Next, we analyzed the sequencing reads covering *CDC42* and *TRAF7* genes in the 5 WDPM cases in this study. Stevers *et al.* reported two unique mutations in *CDC42* and six unique mutations in *TRAF7*. We focused on the corresponding gene regions in the WDPM cases in our study, as summarized in **Supplementary Table 6**.

We identified three unique very low confidence mutations in *TRAF7* gene, one in WDPM-03 and two in WDPM-01 (**Supplementary Table 6**). In WDPM-03, only 16 reads (out of 111 reads) supported the $TRAF7^{Y621D}$ mutant allele. In WDPM-01, $TRAF7^{N520SD}$ was supported by 1 read (out of 102 reads) and $TRAF7^{G536S}$ was supported by 5 reads (out of 109 reads). Given that the tumor cellularity of the WDPM tissues were estimated to be about 50%, the *TRAF* mutations mentioned above were deemed very low confidence and hence did not pass our mutation filtering criteria. The rest of the *CDC42* and *TRAF7* mutated regions reported by Stevers *et al.* were identified as wild type in the WDPM cases in this study.

Thus, within the experimental settings of our study, we do not find any high confidence mutations in *TRAF7* or *CDC42*.

### Supplementary Figures

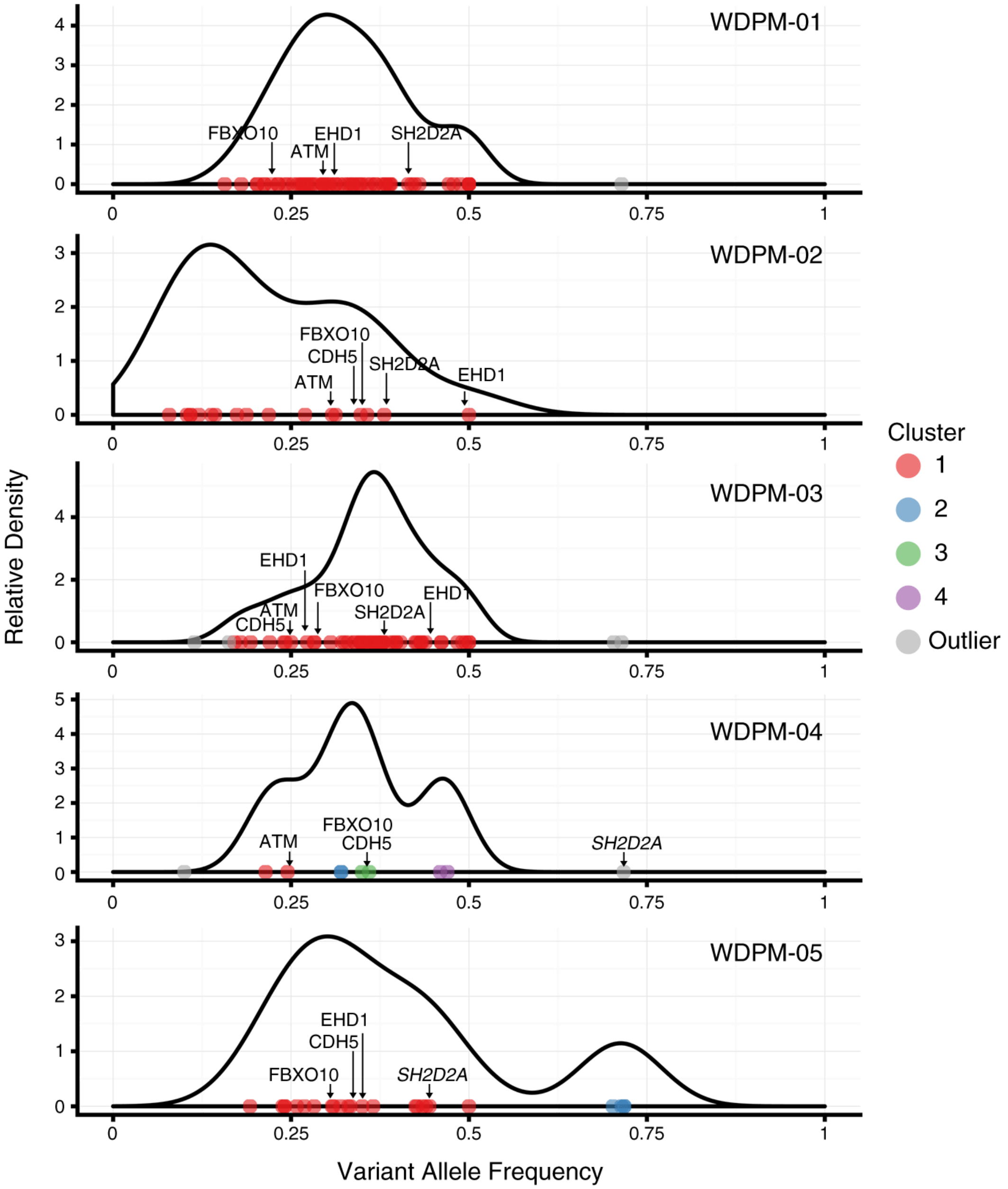


**Supplementary Figure 1**. Distribution of variant allele frequency (VAF) in WDPM. Based on VAF, the somatic mutations identified in WDPM were clustered into different groups using the R-package Maftools.

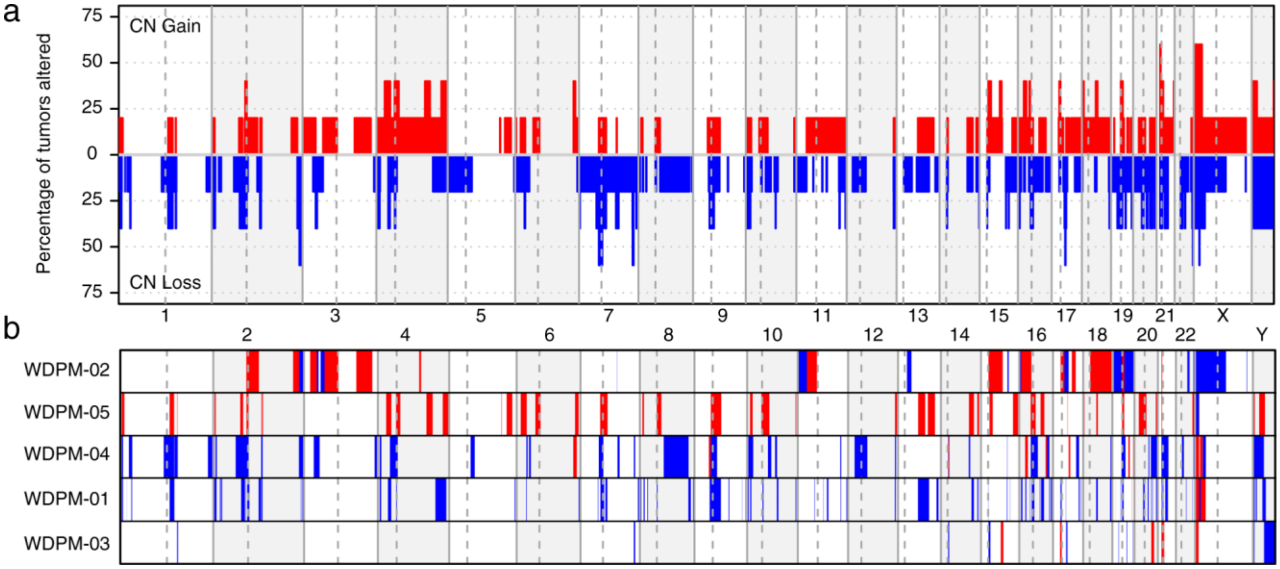


**Supplementary Figure 2.** Landscape of copy number alterations in WDPM. The red and blue color represents the copy-number gains and copy-number loss respectively. (a) Aggregate copy-number alterations by chromosome regions in WDPM. (b) Sample-wise view of copy-number alterations.

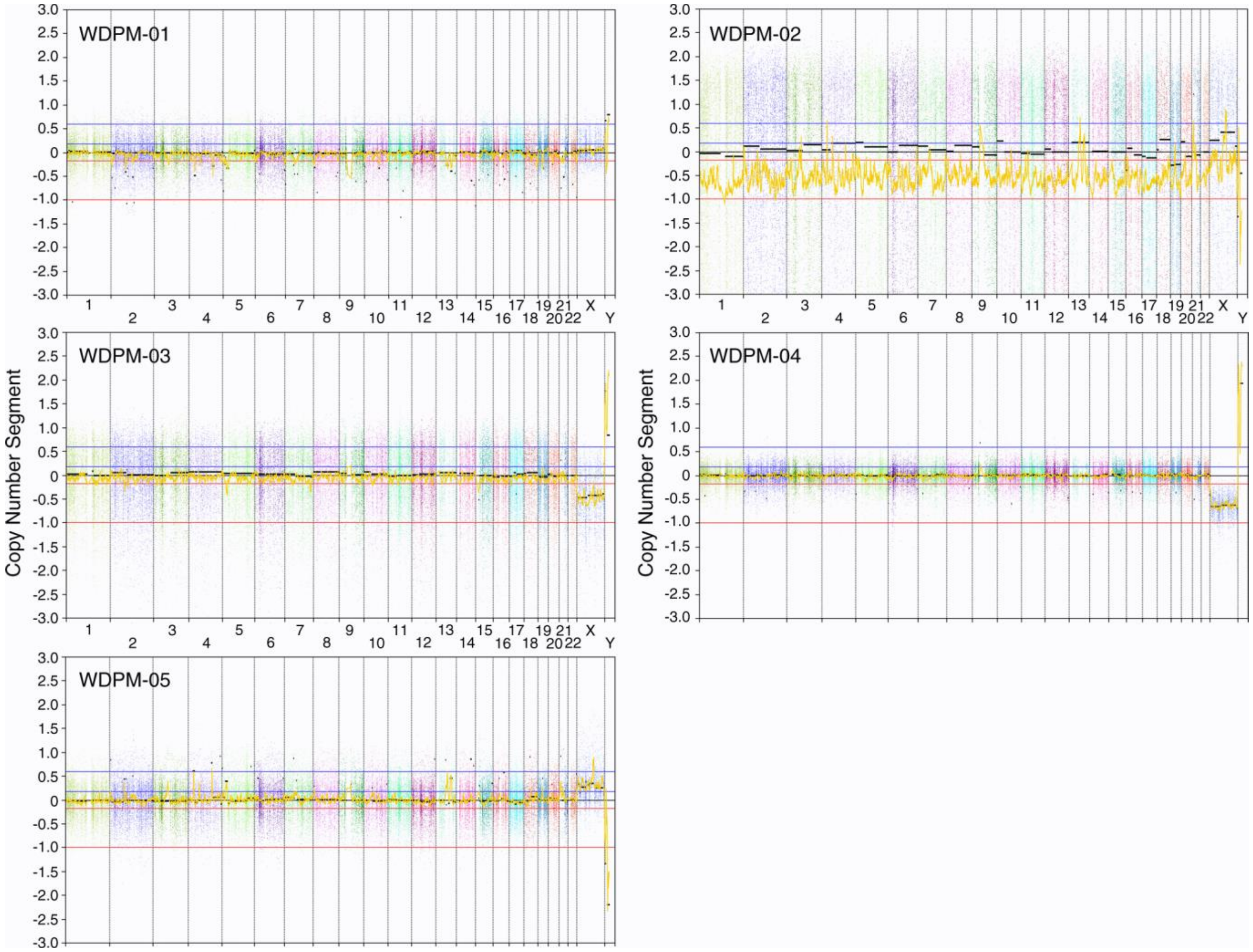


**Supplementary Figure 3**. Copy number segments (log ratio) of WDPM samples.

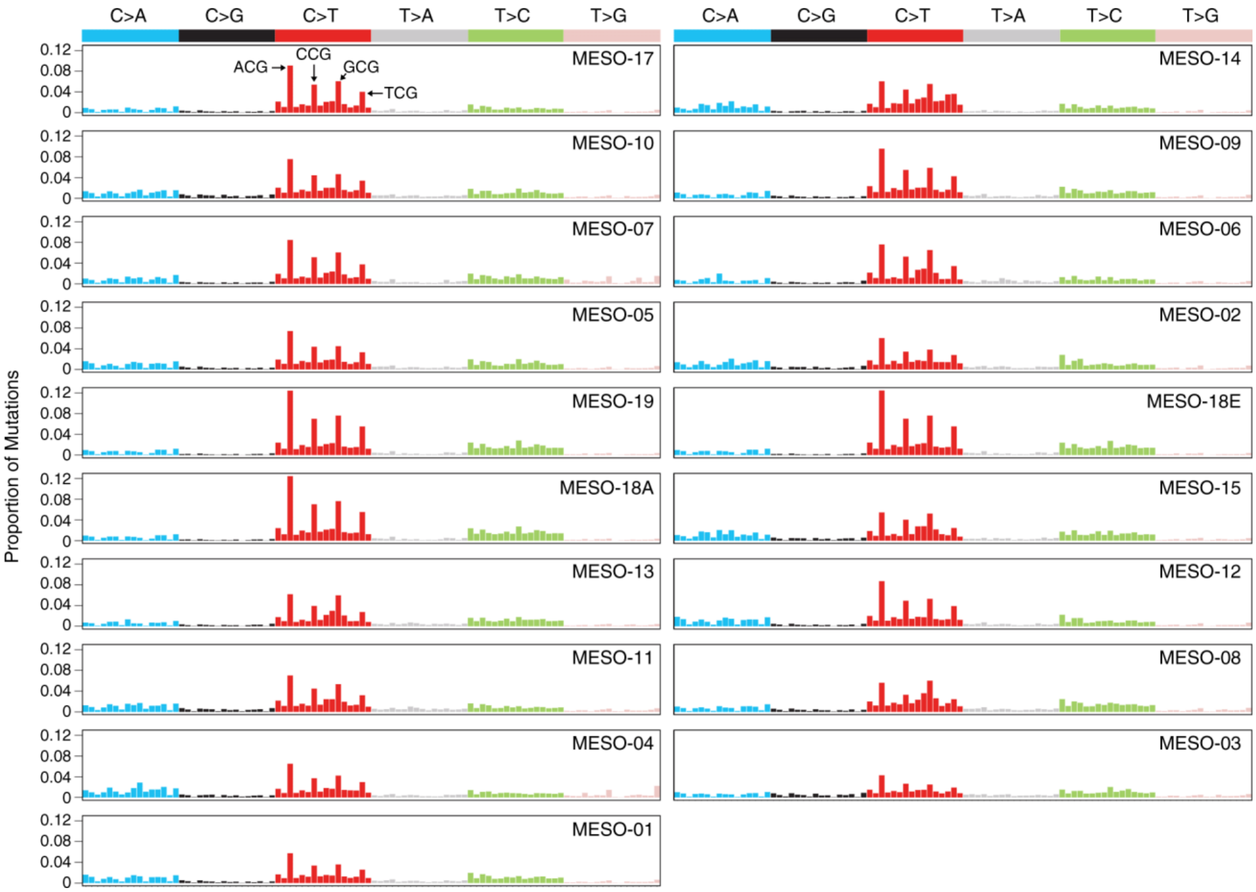


**Supplementary Figure 4**. Mutational signature present in malignant peritoneal mesothelioma obtained from Shrestha *et al*, Genome Medicine 2019.

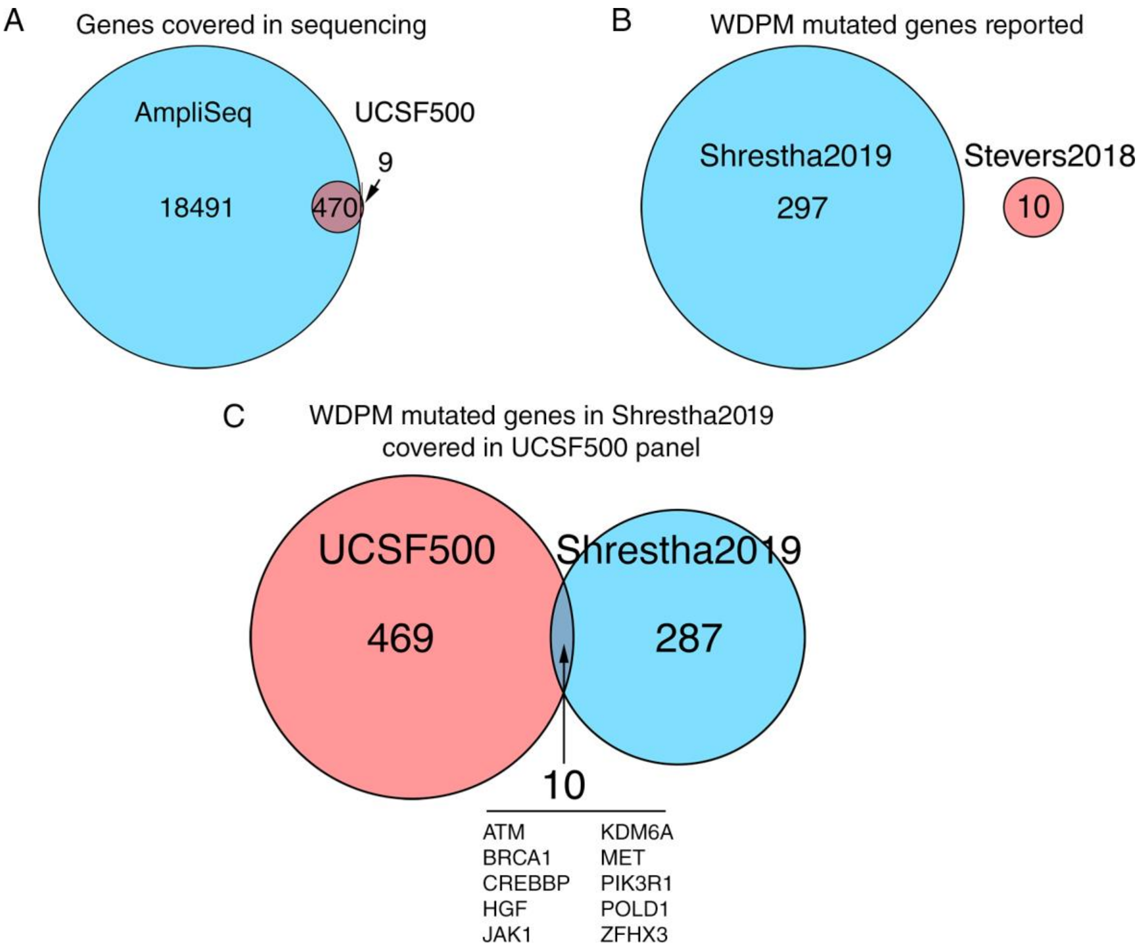


**Supplementary Figure 5. Comparison of Stevers *et al.* 2018 with Shrestha *et. al.* 2019 (present study).** (A) Venn diagram of sequencing target coverage between AmpliSeq Exome Sequencing (used by Shrestha2019) and UCSF500 gene panel (used by Stevers2018). (B) Venn diagram of WDPM mutated genes reported in Stevers2018 and Shrestha2019. (C) Venn diagram of WDPM mutated genes in Shrestha2019 covered in UCSF500 gene panel.
